# Neuropathology and cognitive performance in centenarians

**DOI:** 10.1101/298935

**Authors:** Andrea B. Ganz, Nina Beker, Marc Hulsman, Sietske Sikkes, NBB, Philip Scheltens, August B. Smit, Annemieke J. M. Rozemuller, Jeroen J.M. Hoozemans, Henne Holstege

## Abstract

With aging, the incidence of neuropathological hallmarks of neurodegenerative diseases increases in the brains of cognitively healthy individuals. It is currently unclear to what extent these hallmarks associate with symptoms of disease at extreme ages. Forty centenarians from the 100-plus Study cohort agreed to post-mortem brain donation. Centenarians self-reported to be cognitive healthy at baseline, which was confirmed by a proxy. Objective ante-mortem measurements of cognitive performance were associated with the prevalence, distribution and quantity of age- and AD-related neuropathological hallmarks. Despite self-reported cognitive health, objective neuropsychological testing suggested varying levels of ante-mortem cognitive functioning. Post-mortem, we found that most neuropathological hallmarks related to age and neurodegenerative diseases such as Aβ and Tau pathology, as well as atherosclerosis, were abundantly present in most or all centenarians, whereas Lewy body and pTDP-43 pathology were scarce. We observed that increased pathology loads correlated across pathology subtypes, and an overall trend of higher pathology loads to associate with a lower cognitive test performance. This trend was carried especially by the presence of neurofibrillary tangles (NFTs) and granulovacuolar degeneration (GVD) and to a lesser extent by Aβ-associated pathologies. Cerebral Amyloid Angiopathy (CAA) specifically associated with lower executive functioning in the centenarians. In conclusion, we find that while the centenarians in this cohort escaped or delayed cognitive impairment until extreme ages, their brains reveal varying levels of disease-associated neuropathological hallmarks, some of which associate with cognitive performance.

## Introduction

The increase in average life expectancy warrants gaining insight in healthy aging. In particular, the study of centenarians might reveal new leads towards healthy aging and potential clues to overcome age-related diseases. Alzheimer’s disease (AD) is the most common age-related neurodegenerative disease for which increasing age is the most important risk factor. The prevalence of AD is estimated to double every 5 years after the age of 65, resulting in a population prevalence of roughly 40% for western Europeans aged 90 years and older [22, 28, 32]. The two most prominent neuropathological hallmarks of AD are (1) the intracellular accumulation of hyperphosphorylated Tau protein (pTau) in the form of Neurofibrillary tangles (NFTs) and (2) the accumulation of extracellular Amyloid-β (Aβ) fibrils into senile plaques [24, 30]. Additional AD-associated neuropathological changes are: neuritic plaques (NPs), where Aβ plaques and dystrophic neurites occur in the same structures, cerebral amyloid angiopathy (CAA) illustrated by the deposition of Aβ in the endothelial cells of blood vessels, and granulovacuolar degeneration (GVD), which is characterized by the accumulation of stress proteins in intracellular granules. Whereas Lewy body (LB) and phosphorylated TDP-43 (pTDP-43) proteinopathies are usually associated with other neurodegenerative diseases, they are also observed in late-onset AD cases [38].

Clinicopathologic correlation studies revealed that the severity of cognitive impairment in AD correlates best with the burden of neocortical NFTs [28]. The presence of Aβ is a prerequisite for AD and may catalyze the pathological process that ultimately leads to clinical dementia [38]. However, the concurrent accumulation of individual pathologies, like CAA, GVD, LB, and TDP43, may have an additive effect on the rate of cognitive decline, and contribute to the apparent decrease of the association between NFTs, Aβ and dementia symptoms in older ages [38]. Brains of older dementia patients commonly show less NFT and Aβ pathology compared to younger AD patients [8, 14, 15, 34]. This suggests that, while disease-associated proteinopathies distinguish well between brains from young AD cases and age-matched non-demented controls, this is more complicated for brains from older individuals. So far, post-mortem brain assessments of cognitively healthy individuals who reached >110 years revealed various results: some cognitively healthy centenarians did not accumulate significant pathology, while others appeared to have relatively high levels of pathology [10, 29, 39].

Previous studies on brain tissues from centenarians commonly lack information on cognitive well-being shortly before death, which is essential for studying the association with neuropathological hallmarks in post mortem brain tissue.

In this study, we report on the occurrence of neuropathological changes in a unique, post mortem brain collection of self-reported cognitively healthy centenarians from the 100-plus Study cohort [16]. The aim of this study is to investigate how neuropathological changes relate with performance on neuropsychological tests in centenarians. Prospective neuropsychological testing of these individuals allowed us to correlate ante-mortem cognitive functioning with the occurrence of pathological hallmarks of neurodegenerative diseases in the post-mortem brain. For this, we focused on inclusions of pTau as NFTs [4] and ARTAG [21]; the distribution of Aβ protein as diffuse [41], classical or NPs [23], and CAA [40]; the distribution of casein kinase 1 delta as a marker of GVD [43], and the distribution Lewy bodies [6] and pTDP-43 accumulations [25]. Furthermore, for an individual case, we report on the in-vivo assessment of pathology using an MRI and amyloid PET scan, and the occurrence of post-mortem neuropathological changes.

## Material and methods

### 100-plus Study cohort

The 100-plus Study is a prospective centenarian cohort study. Inclusion criteria for the study are, next to being able to provide proof of being 100 years or older, that centenarians “self-report to be cognitively healthy, which is confirmed by a proxy” [16]. Although it is not a requirement for study participation, almost 30% of all participants agree to post-mortem brain donation. Here we compare the ante- and post-mortem status of the first 40 participants of the 100-plus Study that came to autopsy.

The 100-Plus Study was approved by the Medical Ethical Committee of the VU University Medical Center. Informed consent was obtained from all study participants. The detailed study protocol is described elsewhere [16].

### Post-mortem brain autopsy procedures

Autopsies were performed in collaboration with the Netherlands Brain Bank (NBB, Amsterdam, The Netherlands). Brain weight was recorded, and we macroscopically determined levels of atrophy and atherosclerosis of the intracranial vessels. Atrophy was subjectively staged by an experienced neuropathologist according to severity (none (0), slight (1), moderate (2) or strong (3)). Atherosclerosis was scored as mild (1), if only some parts of the circle of Willis and the basal arteries were affected; moderate (2), if the circle of Willis and the basal arteries were severely affected and the arteria cerebri media was not affected beyond the first cm; and severe (3), if also the deeper part of the arteria cerebri media was affected. The right hemisphere of the brain was formalin fixed, and the left hemisphere was dissected and the pieces were snap frozen in liquid nitrogen.

### Neuropathology: immunohistochemistry

Neuropathological characterization of centenarian brains was performed by Haematoxylin and Eosin (H&E) stain, Gallyas silver stain [31, 44] and immunohistochemistry (IHC). For IHC, brain tissues were formalin fixed and paraffin embedded (FFPE), then sectioned into 6 μm slices and mounted on microscope slides (Leica Xtra adhesive slides, Leica Microsystems, Rijswijk, The Netherlands). After deparaffinization and rehydration, endogenous peroxidase activity was blocked in 0.3% H_2_O_2_ in methanol for 30 minutes. Antigen retrieval was performed with heated sodium-citrate buffer (10 mM/L, pH 6.0). For the staining of α-synuclein, pretreatment included only heated sodium citrate without endogenous peroxidase blocking. All primary antibodies were diluted in normal antibody diluent (Immunologic, VWR company, Duiven, The Netherlands) (Table 1). Primary antibody incubation was performed overnight at 4°C. Subsequently, sections were incubated with the EnVision detection system (goat anti-mouse/rabbit horseradish peroxidase (HRP), DAKO, Heverlee, Belgium) for 40 minutes at room temperature. Between incubation steps, sections were rinsed with phosphate buffered saline, pH 7.0 (PBS). Sections were incubated for 5 minutes with the chromogen 3,3’-diaminobenzidine (DAB, EnVision Detection system/HRP, DAKO, Heverlee, Belgium) to visualize immunoreactivity. Nuclei were counterstained with haematoxylin. Hereafter, slides were dehydrated and mounted using the non-aqueous mounting medium Quick-D (Klinipath, Duiven, The Netherlands). Negative controls were obtained by performing the same procedures while omitting the primary antibody.

**Table 1:** Overview of primary antibodies.

| Antibody | Antigen | Species | Dilution<br>(FFPE<br>material) | Manufacturer |
| --- | --- | --- | --- | --- |
| **IC-16** | N-terminal amino acids 1-6 of the Aβ peptide | mouse | 1:800 | Kind gift of Prof. Dr. Korth, Heinrich Heine University, Düsseldorf, Germany [46] |
| **AT-8** | Tau phosphorylated at Ser202 and Thr205 | mouse | 1:800 | Pierce Biotechnology, Rockford, IL, USA |
| **CK1δ** | casein kinase 1 delta | mouse | 1:100 | Santa Cruz Biotechnology, Heidelberg, Germany |
| **pTDP-43** | Transactive response DNA binding protein 43 | mouse | 1:8000 | Cosmo Bio, Tokyo, Japan |
| **18-0215** | alpha-synuclein | mouse | 1:200 | Zymed, Thermo Fisher Scientific, Bleiswijk, The Netherlands |

### Neuropathological evaluation

Distribution of Aβ and hyperphosphorylated tau (pTau) in NFTs and NPs was determined by immunohistochemical staining (IHC) with respectively IC-16 and AT-8 antibodies on the frontal cortex (F2), temporal pole cortex, parietal cortex (superior and inferior lobule), occipital pole cortex and the hippocampus (CA1 and entorhinal area of the parahippocampal gyrus). NFTs and NPs were identified using a combination of Gallyas silver staining and IHC for pTau (AT-8). Aβ distribution was evaluated according to the Thal staging ranging from 0 to 5 [42]. The temporal-spatial distribution of NFTs was evaluated according to the Braak stages ranging from 0 to VI [3–5]. NPs were evaluated according to CERAD scores ranging from 0 to 3 [23]. The presence of ARTAG was assessed using IHC for pTau in the frontal cortex (F2), temporal pole cortex, parietal cortex (superior and inferior lobule), occipital pole cortex and the hippocampus (CA1 and entorhinal area of the parahippocampal gyrus).

GVD was assessed by IHC using CK1δ as marker, and scored based on the staging system from Thal et al., 2011 [43], ranging from 0 to 5. The presence of CK1δ positive accumulations was assessed in the hippocampal areas CA1/Subiculum and CA4, entorhinal cortex, temporal pole cortex, amygdala, cingulate gyrus, frontal (F2) and parietal lobe. GVD stage classified as 1, if only the subiculum and CA1 region of the hippocampal formation were affected. In stage 2, the CA4 and/or entorhinal cortex were additionally affected. Additional CK1δ granules in the temporal pole cortex classified stage 3. Cases were scored as stage 4 if the amygdala was affected, but neither the cingulate gyrus or the frontal or parietal cortex were affected. If either cingulate gyrus, frontal cortex or parietal cortex showed additional depositions of CK1δ, cases were classified as stage 5.

pTDP-43 pathology was assessed by IHC and stages ranged from 0 to 3 according to Nag et al., 2015 [25]. Immunohistochemistry with the pTDP-43 antibody was performed on slices of the amygdala (stage 1), hippocampal areas CA1/Subiculum and dentate gyrus, entorhinal cortex (stage 2), frontal and temporal pole cortex (stage 3), in which the presence of cytoplasmic inclusions was assessed.

Lewy body pathology was assessed according to Braak [2, 6] by IHC for α-synuclein in the hippocampus, amygdala, medulla oblongata, substantia nigra and locus coeruleus. For a case with high amounts of α-synuclein inclusions, the frontal and precentral gyrus were additionally assessed.

### Neuropsychological tests and educational attainment

All study participants were visited at home by a trained researcher at baseline inclusion, and when available and willing, at yearly follow-up visits. During these visits, we objectively assessed cognitive functioning using an elaborate test battery described elsewhere [16]. Here we report the performance on the Mini-Mental State Examination (MMSE), Visual Association Test (VAT) and the Clock Drawing Test (CDT). The MMSE is an 11-item test evaluating overall cognitive functioning with a scoring range of 0-30 (bad-good) [13]; missing items were imputed as described before [16], provided no more than 6 items were missing. The VAT is a test for episodic memory with a 0-12 (trial 1+2) range (bad-good) [17], and the CDT is a test for visuospatial and executive functioning, with a 0-5 range (bad-good) [35, 36]. When sensory impairments, fatigue, or lack of motivation impeded test administration, no score was reported (N/A). We report test scores at baseline and at last visit before death. For cases with only one visit, baseline- and last visit are the same. Number of visits and time between last visit and death is listed in Supplementary Table 2. Educational attainment was assessed by self-report and categorized according to the International Standard Classification of Education 1997 (ISCED)[45], as well as years of education.

### Correlation of cognitive performance with post mortem brain pathology

To determine whether performance on cognitive tests of centenarians correlated with postmortem brain pathologies, we correlated pre-mortem neuropsychological test scores and demographic descriptives with levels of post-mortem neuropathological hallmarks of aging and disease For this, we used MMSE, VAT and CDT test scores, both at baseline and last visit. A secondary measure of cognitive performance was “the ability to complete all tests” as a proxy of fatigue or non-willingness; scores were 2 (all tests completed, or tests not applied due to sensory problems), 1 (MMSE completed, which was administered first, but not VAT and/or CDT), and 0 (no tests completed). After each study visit, the impression of cognitive health of the participant was subjectively estimated by a trained researcher as 0 (symptoms of impairment), 1 (possible symptoms of impairment), or 2 (no symptoms of impairment) (see [16] for protocol). The scored impression of cognitive health closest to death was used for analysis. ApoE genotypes were ranked according to their association with increased AD-risk (odds ratios) as reported previously [7]: E2/2 (OR 0.24); E2/3 (OR 0.5); E3/3 (OR 1.0); E2/4 (OR 3.2); E3/4 (OR 5.5), E4/4 (OR 20.6). Education was categorized by years of formal education. Sex: “being male” (female, 0; male, 1). As post-mortem characteristics of all 40 brains, we used brain weights corrected for sex by linear regression and the levels of atrophy, atherosclerosis, scored as described above. For a subset of 35 brains Thal stage GVD, Infarcts, pTDP-43 staging was available, and for 26 brains Braak stage for Lewy bodies, Thal stage for CAA, Thal stage for Aβ, CERAD score for NPs, Braak stage for NFT distribution and hippocampal sclerosis was available. Overview of all variables is provided in Supplementary Tables S1 and S2.

### Statistical analyses

Correlations between neuropsychological test performance and neuropathology were determined using Pearson correlation. We calculated p-values and false discovery rates (fdr) for all correlations (Supplemental Tables 3b and 3c). In Figure 3, strength of the Pearson correlation coefficient is indicated by the size and color of each dot, the fdr is indicated as asterisks (^∗^<20%, ^∗∗^<10%, ^∗∗∗^<5%). All calculations were performed in R (version 3.3) [33].

### PET/MRI scan in case report

Scanning was performed 1 month after study inclusion for case 2015-060. The participant was injected with 383 MBq 11C-PIB. Sixty minutes after injection, dynamic images of the brain were captured on the Philips TF PET-MRI scanner in 6 frames of 5 minutes duration. Due to extensive movement, only the activity in the first frame could be used. The PET scan was visually assessed by an experienced nuclear medicine physician.

## Results

We analyzed brain tissue from 40 centenarians, (72.5% female), aged between 100 and 111 years (Supplementary Table S2).

### Neuropathological characteristics in centenarian brains

The mean post-mortem interval for all 40 donated brains was 6h44min (range: 3.5 - 12 hours). At autopsy the median brain weight was 1195 g (IQR 1060 g-1355 g) in males and 1115g (IQR 965 g-1320 g) in females. ApoE genotypes were: E2/3 (15.8%), E3/3 (73.7%), E2/4 (5.3%), and E3/4 (7.9%). Full neuropathological characterization was present for 26 brains, partial characterization with staging for pTDP-43 stage and Thal stage GVD^∗^ was present for 35 cases. We observed varying levels of atherosclerosis in all 40 brains. Mild or moderate atrophy was present in 50% percent of all cases, while severe atrophy was not observed. All 26 centenarian-brains with full neuropathological assessment revealed known age-related pathologies such as ARTAG [21] (Figure 1C and Supplementary Table 2), and were scored with high stages of GVD: only GVD stages between 3 and 5 were observed (Figure 1I). According to the NIA-AA guidelines, 8% of the centenarians had no AD-associated neuropathological changes, 42% had low neuropathological changes, 50% had intermediate changes, and none had high level of changes (Figure 1G). Levels of Aβ deposits cover all possible Thal stages (mean Thal stage for Aβ: 2.6). The NFT distribution ranged from Braak stage I to IV, (mean Braak stage: 3). None of the centenarians had Braak stage 0 or Braak stage V or VI. The level of NPs ranged from CERAD scores 0 to 2, CERAD score 3 was not observed (mean CERAD score: 0.8). In 20 of the 26 centenarians we observed CAA pathology (77%, mean Thal stage for CAA: 1), always in combination with Aβ plaques. We observed pTDP-43 pathology in 13 of 35 centenarians (37%, mean pTDP-43 stage: 0.9), and 5 cases show additional hippocampal sclerosis (19.2%). Hippocampal sclerosis was exclusively observed in combination with pTDP-43 pathology (Supplementary Table 2). Lewy body pathology was observed in 4 of 26 cases (15.4%, mean Braak stage for Lewy bodies: 0.4), 3 of which co-occurred with pTDP-43 pathology. Despite high cognitive test performance (baseline score on the MMSE: 27), one case was clinically diagnosed with Parkinson’s disease many years before entering the study, the brain included many Lewy bodies at post-mortem neuropathological evaluation (Braak stage for Lewy bodies VI). Additionally, we detected infarcts in 14 (54%) of the 26 centenarians.

**Figure 1:**
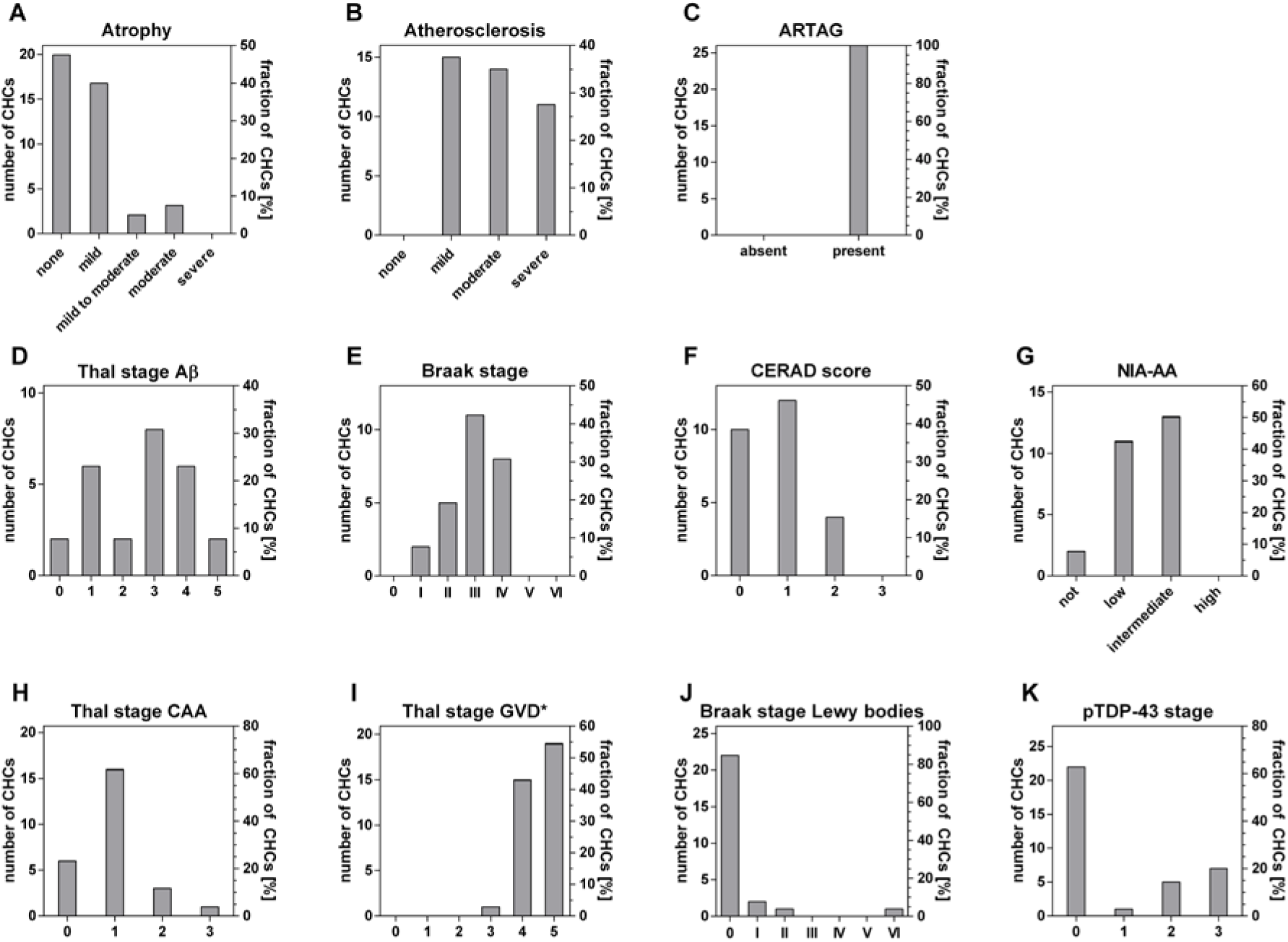
Distribution and severities of pathology within the group of 26 centenarians. For atrophy and atherosclerosis, n=40. n=35 for Thal stage GVD^∗^ and pTDP-43 stage, for all other pathologies, n=26. Thal stage GVD^∗^ indicates an adapted staging system, see Methods section.

Overall, the centenarians in our cohort showed moderate levels of AD related pathology. Centenarians rarely showed the highest score for AD pathology, with the exception of GVD. Overall scores for Lewy bodies and TDP-43 appeared relatively low in centenarians.

### Performance on neuropsychological tests of 40 centenarian brain donors

The aim of this study is to investigate how neuropathological changes are associated with performance on neuropsychological tests in centenarians. The last visit occurred on average 10±7 months prior to brain donation. 18 centenarians were available for at least one follow-up visit; follow-up for two additional centenarians was performed by questionnaire. Centenarians had a variable educational attainment (median years of education: 8, range 6 - 20y). MMSE, VAT and CDT scores were available for respectively 95%, 67.5% and 65% of centenarians at baseline (Figure 2A). At last visit, MMSE, VAT and CDT scores were available for respectively 87.5%, 57.5% and 52.5% of centenarians. The mean (±SD) score on the **MMSE** was 24±4.5 at baseline and 23.7±4.6 at last visit. Mean scores for the **VAT** were 7.7±3.2 at baseline and 6.9±3.6 at last visit respectively (Figure 2B), while **CDT** scores were on average 2.9±1.3 at baseline and 3.3±1.5 at last visit (Figure 2C). Detailed demographics, neuropsychological and neuropathological measurements at the individual level are listed in Supplementary Table S2.

**Figure 2:**
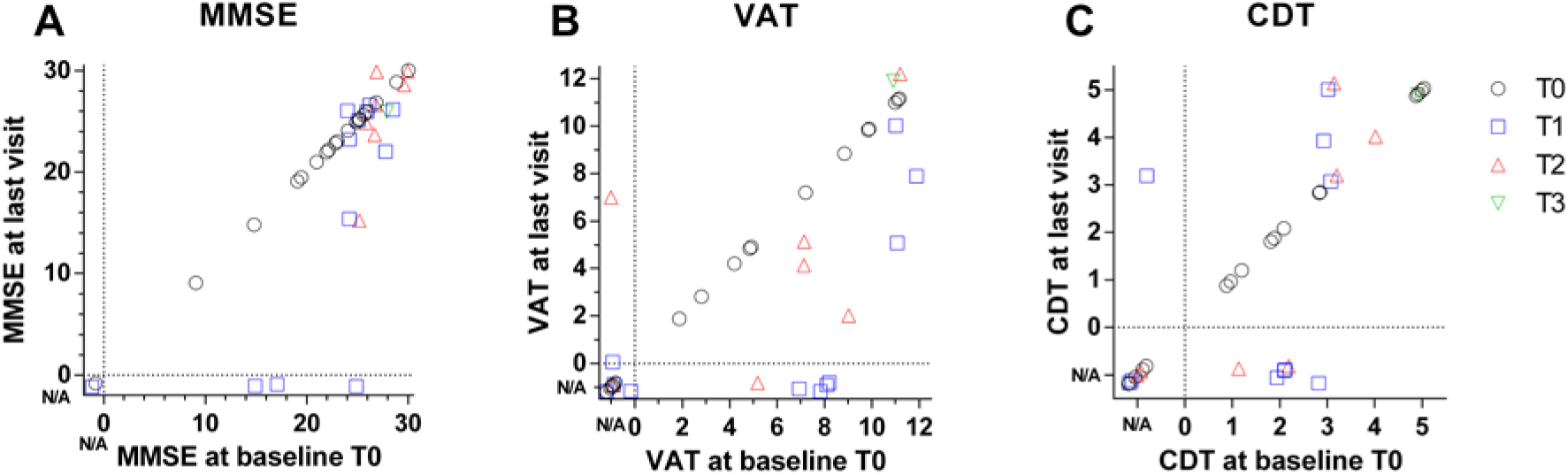
Distribution of performance on selected neuropsychological tests within the group of centenarians at baseline and last visit. For all neuropsychological tests, n=40. If a test was not administered or not completed, the score was indicated as not available (N/A), set apart from the scores by a dashed line. Dots are scattered to ensure visibility of all points. Cases for which only baseline data is available (T0) are shown as black circles, cases with one (T1), two (T2) or three (T3) follow-up visit are shown as blue squares and red or green triangles respectively.

### Correlations between different neuropathological hallmarks

The levels of neuropathological hallmarks of AD are inter-correlated (Figure 3 and Supplementary Table 2). Particularly, Braak stage for NFTs correlated significantly with GVD levels (false discovery rate (fdr=0.002), and to a lesser extent with CERAD scores (fdr=0.196). Thal stage for Aβ correlated significantly with levels of CAA and Braak stages (fdr=0.8; and fdr=0.16 respectively), and as expected with CERAD scores, of which it is a part (fdr=4.46E-16). TDP-43 stage is strongly correlated with hippocampal sclerosis (fdr=0.044). In contrast, the levels of alpha-synuclein were not associated with the levels of any other assessed neuropathology.

**Figure 3:**
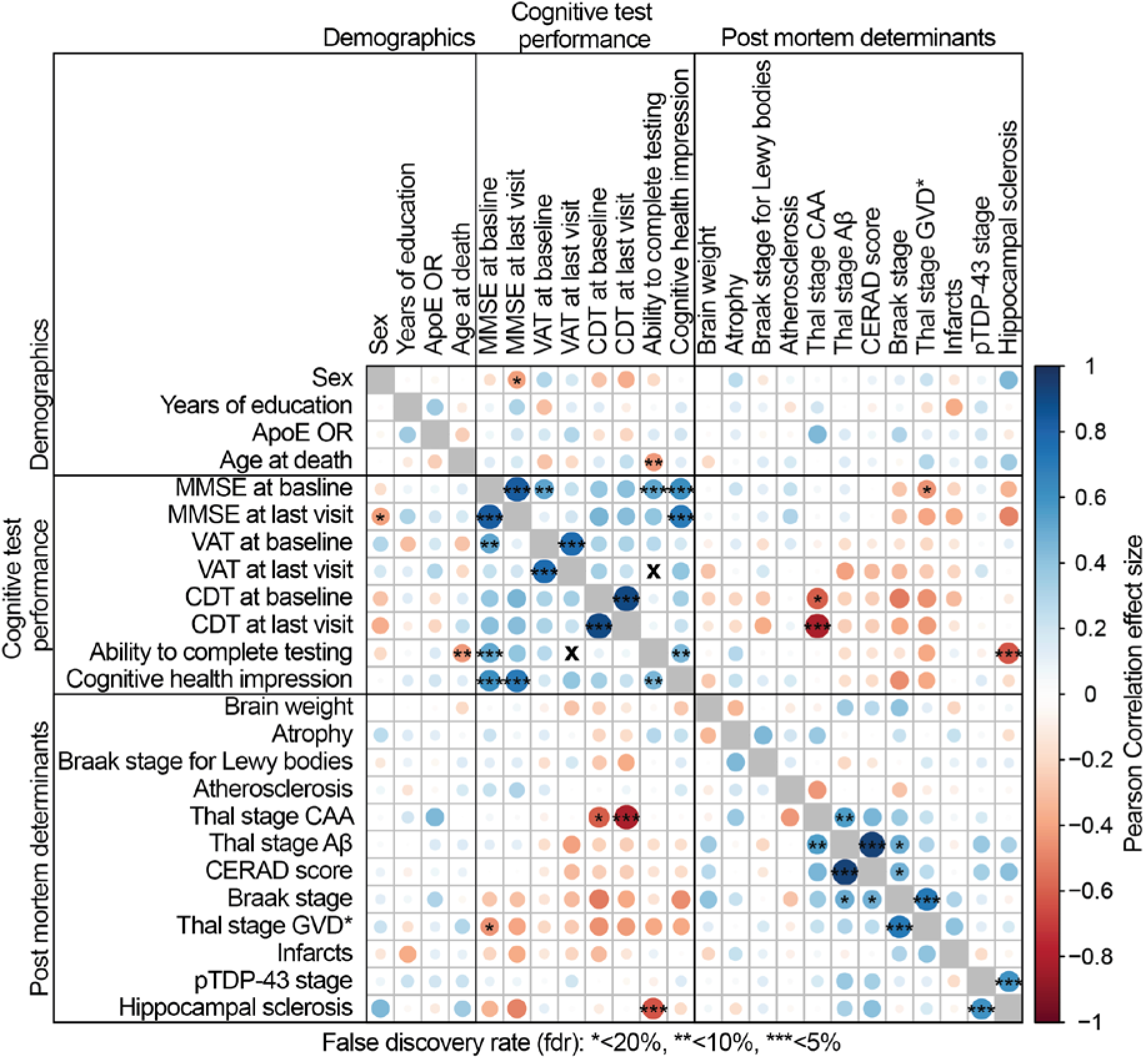
Pearson correlation plot with false discovery rate (fdr). Color and size of the circles indicate the strength of the Pearson correlation coefficient (for Pearson correlation coefficient, p-value and numerical fdr see Supplementary Tables 3a-c), asterisks indicate the false discovery rate (^∗^<20%, ^∗∗^<10%, ^∗∗∗^<5%). Correlations between the same test at baseline and last visit were performed for 20 cases with at least one follow up visit. Thal stage GVD^∗^ indicates that we utilized a staging system for GVD adapted from Thal et al., 2011[43].

### Correlations between test performance, demographic characteristics and pathology

We observed an overall trend that better performance correlates among cognitive tests. For the centenarians who completed cognitive tests at baseline and at last visit, we find a strong correlation between performances (Figure 3). The ability to complete testing was associated with lower ages at death, researcher impression of cognitive health and higher baseline MMSE scores. Likewise, we observed that increased pathology loads correlate among pathology subtypes. We also observe an overall trend that lower pathology loads correlate with better performance on tests (Figure 3). This trend is mainly carried by the association of Braak stages for NFTs and higher GVD stages with lower cognitive performance across all tests. Higher Thal stages for Aβ pathology and higher CERAD scores also show this trend with lower test performance, but to a much lesser extent. Two specific associations stand out: increased loads of CAA correlated significantly with a low CDT performance at baseline (fdr=0.156) and, despite a smaller group-size, this association was intensified for CDT performance at last visit (fdr=0.02). Secondly, the presence and severity of hippocampal sclerosis correlated with a decreased ability to complete testing at follow up visits (fdr=0.03) and non-significantly with a lower performance on the MMSE, both at baseline (fdr=0.46), and last visit (fdr=0.24). GVD stage only correlated with a low MMSE performance at baseline (fdr=0.16). This correlation was lost at last visit (fdr=0.25).

### Case report of 103 year old female

Case 2015-060 was a female centenarian with a fit physique included in the 100-plus Study at the age of 102. She finished first stage of tertiary education, and had enjoyed a total of 18 years of education (ISCED level 5). At baseline she scored 27 points on the MMSE. She volunteered to undergo an MRI and PIB-PET scan one month after inclusion into the study. The MRI scan revealed moderate hippocampal atrophy (MTA score 2), white matter abnormalities (Fazekas score II), and few microbleeds. The PET scan using [^11^C]PiB as an amyloid tracer was positive throughout the brain, with the highest signal in the frontal area (Figure 4). The centenarian died 10 months after inclusion of cachexia. Proxies reported maintained cognitive health until the terminal phase. Brain autopsy was performed with a post mortem delay of 8.5 hours. Overall, no major abnormalities were observed at the macroscopic level. Slight atrophy of the temporal lobe was observed and arteries were slightly atherosclerotic. Two small meningeomas were observed in the left part of the scull, as well as a small (0.6 cm) old cortical infarct in the medial frontal gyrus. The substantia nigra appeared relatively pale. Neuropathological evaluation revealed intermediate AD neuropathologic changes (A3B2C1). Consistent with the positive amyloid PET scan in vivo, we observed Aβ pathology in the form of classical and diffuse plaques (Thal stage Aβ 4), as well as capillary CAA-type 1 (capCAA) (Thal stage CAA 2) (Figure 5 panel d-f) during the post-mortem neuropathological analysis. Diffuse Aβ deposits were detected in the frontal, frontobasal, occipital, temporal and parietal cortex, the molecular layer, CA1 and subiculum of the hippocampus, as well as the caudate nucleus, putamen and nucleus accumbens. A relatively low number of classic senile plaques with a dense core was observed throughout the neocortex. Prominent presence of CAA was observed in the parietal cortex and the leptomeninges. Capillary CAA (type 1) was observed in moderate levels in the frontal cortex (Figure 5, panel f), the cerebellum, sporadically in the occipital cortex, and was absent in the temporal cortex. Together, the relatively high levels of CAA might explain the high signal of the [^11^C]PiB-PET, as CAA is also known to be detected by this tracer (Farid et al., 2017[12]), as well as the abnormalities observed on the MRI.

**Fig. 4:**
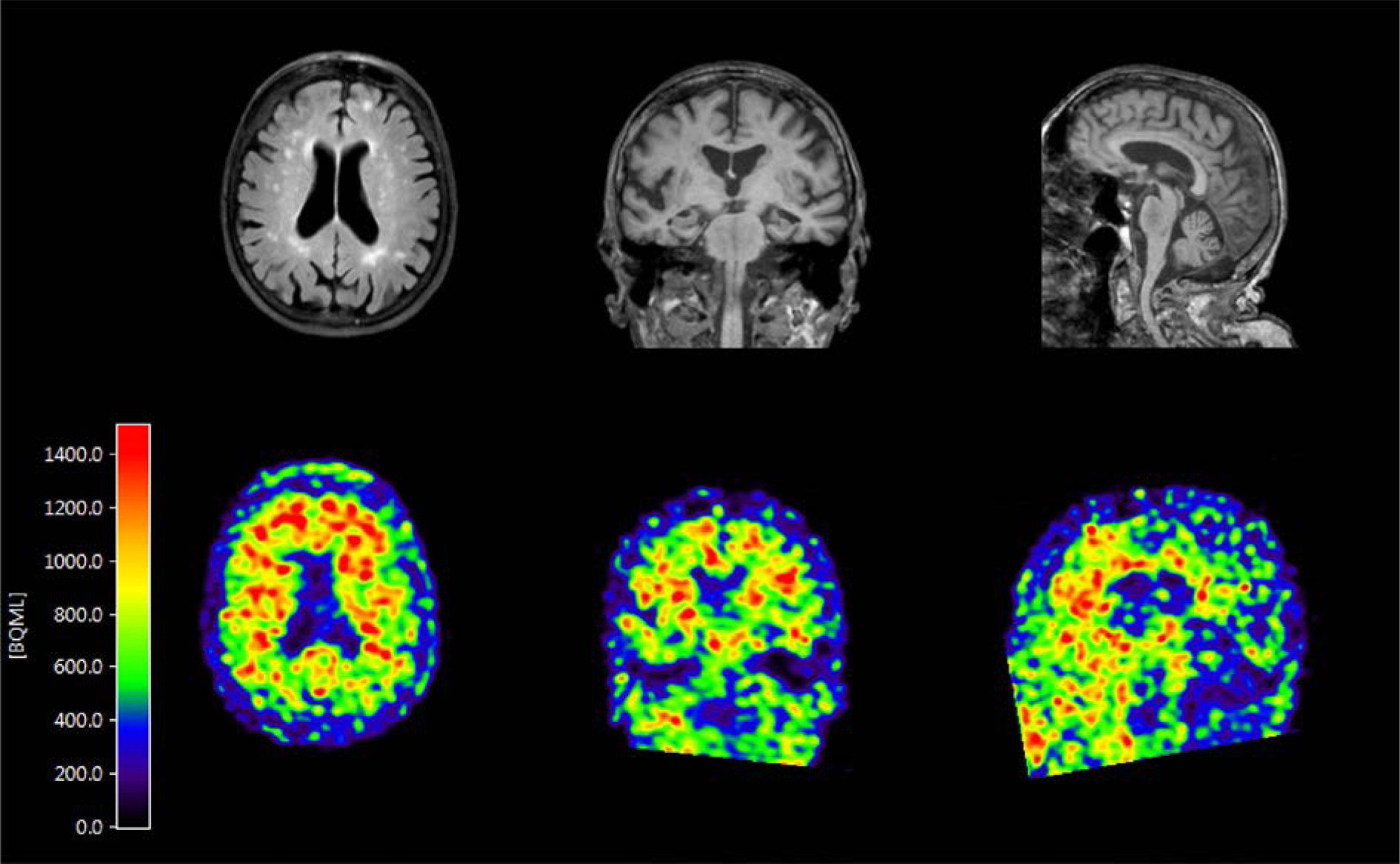
Dynamic [^11^C]PiB-PET and MRI scan for 11C-PIB of case 2015-060. The scan was performed by the ECAT EXACT HR1 scanner (Siemens/CTI) one month after study inclusion. PET scan for amyloid is positive in the frontal area, MRI shows moderate hippocampal atrophy (MTA score 2) as well as vascular lesions in the white matter (Fazekas score II), suggesting amyloid angiopathy.

**Figure 5:**
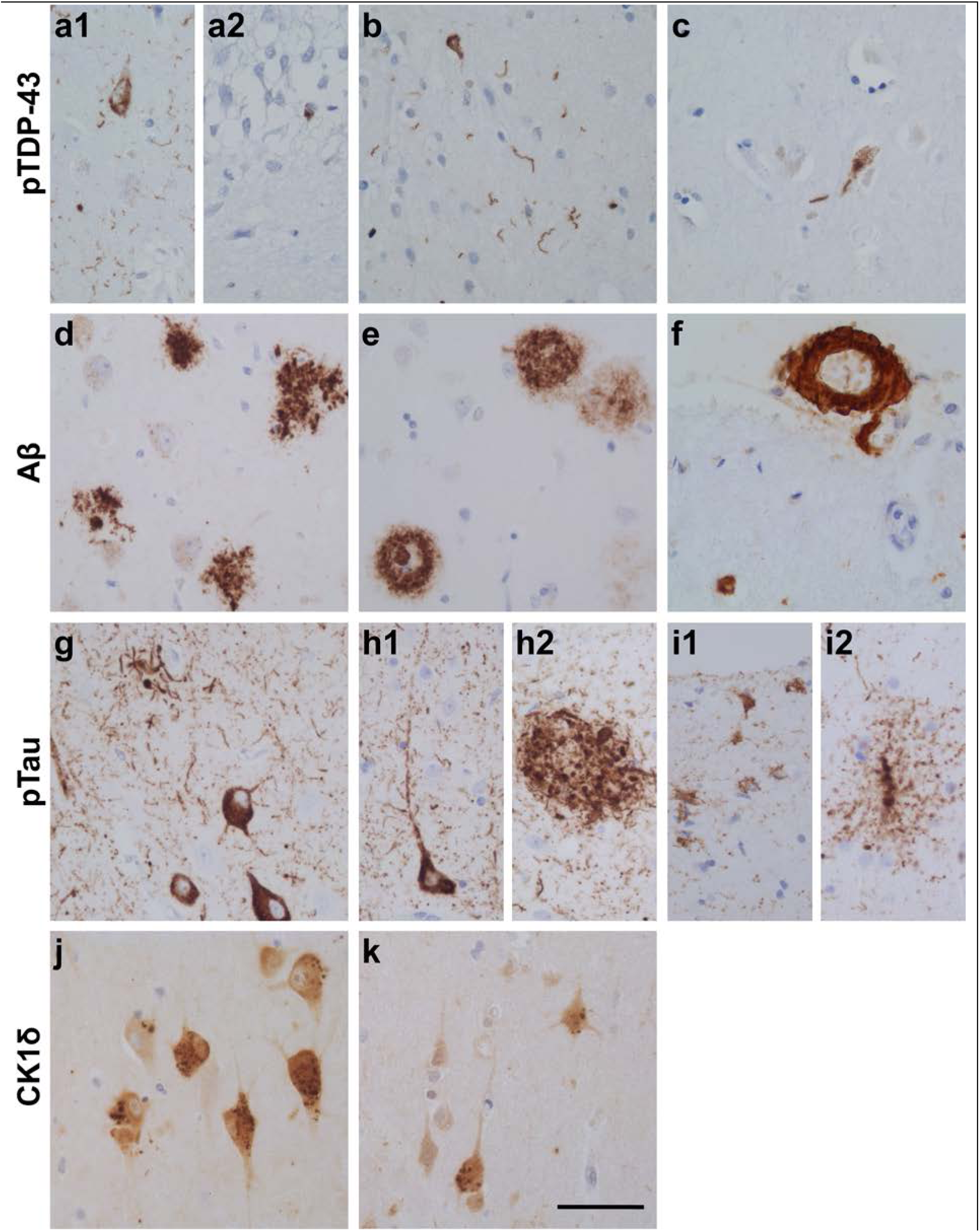
Representative neuropathological lesions found in the hippocampus (a,d,g,j), middle temporal lobe cortex (b,e,h,i,k), amygdala (c) and frontal lobe cortex (f) of case 2015-060. Shown are exemplary pTDP-43 accumulations (a-c) in the hippocampal subregion CA1 (a1) and dentate gyrus (DG) (a2), temporal pole cortex (T) (b) and Amygdala (Amy) (c), Aβ positivity (d-f) in the form of diffuse and classical plaques in the hippocampal CA1 region (d) and temporal pole cortex (e), as well as CAA of large vessels and capillaries in the frontal lobe (F2) (f). pTau immuno-staining (g-i) is shown as (pre)tangles and neuritic plaque like structures in the hippocampal CA1 region (g) and temporal pole cortex (h1 and h2), as well as astroglial pTau in ARTAG in the temporal pole cortex (i1 and i2). GVD (CK1δ granules) (j-k) is shown in the hippocampus (j) and temporal pole cortex (k). Scale bar 25μm.

Immunostaining for pTau showed prominent occurrence of dystrophic neurites, neurofibrillary tangles and pretangles in the hippocampal areas CA1, subiculum and transentorhinal cortex. Moderate numbers of pretangles and threads were observed in the frontal cortex and occipital cortex. Severe presence of pTau immunoreactive neuropil threads and (pre)tangles was observed in the temporal cortex. Additionally, we observed pTau positive glial cells in the temporal cortex and subcortical white matter. In the anterior part of the hippocampus, pTau positive glial cells were observed associated with blood vessels, also indicative of ARTAG [21]. pTau positive neurons and glial cells were also observed in the amygdala and insula. In summary, immuno-positive pTau was observed as NFTs (Braak stage IV) and NP like structures (CERAD score 1), as well as in the form of ARTAG (Figure 5, panel g-i). GVD (CK1δ positive granules) was present in high numbers in the hippocampal subfields CA1 and subiculum, CA4, entorhinal cortex, temporal cortex, amygdala and gyrus cinguli (Thal stage GVD^∗^ 5) (Figure 5, panel j-k). Accumulations of pTDP-43 pathology were widespread, including the amygdala, hippocampus and temporal pole cortex (pTDP-43 stage 3) (Figure 5, panel a-c). No α-synuclein inclusions were observed in hippocampus, substantia nigra, amygdala, locus coeruleus, and in the nuclei of the medulla oblongata. A relative moderate decrease of Purkinje cells was observed in the cerebellum.

## Discussion

In a unique cohort of 40 centenarians, in which individual cognitive functioning was assessed shortly before death, we investigated the post-mortem brain for disease- and aging-associated neuropathological changes. At baseline, cognitive performance varied: neuropsychological test performance suggested that some centenarians had no symptoms of cognitive impairment, while others showed considerable impairment in several cognitive domains. Furthermore, a small subset of centenarians with multiple follow-up visits showed some decline after study inclusion, while others remained cognitively stable until death. We also observe an overall trend that higher pathology loads associate amongst each other, but also with a lower performance on cognitive tests: this trend is carried especially by the NFT and GVD loads and to a lesser extent by Aβ-associated pathologies. Whereas previous findings indicated that many elderly individuals with CAA remain asymptomatic [12], our results suggested that the severity of amyloid accumulation in the vessel walls (CAA), but not Aβ accumulation in the form of plaques, significantly associated with lower executive functioning, as measured by the Clock Drawing Test (CDT). Together, our findings suggest that cognitive performance of centenarians is, at least in part, sensitive to the increase of neuropathological hallmarks of disease.

### Accumulation of neuropathologies is common in centenarians

Aβ pathology was present in most centenarians at variable levels, and Braak stages for NFT were found in all centenarians and distribution occurred up to stage IV, and we did not observe Braak stages 0, V or VI. Similarly, we detected significant accumulations of NPs, which did not exceed CERAD score 2. In fact, several centenarians have exceptionally good brain function with a high NFT load (Braak stage IV) and NP accumulations (CERAD scores 1 or 2), and high Aβ load (Thal stages 3-5). This is intriguing, since most younger individuals with similar amounts of pathology have at least some symptoms of AD [28, 38]. Overall, the cognitive function of these centenarians was conserved despite the presence of neuropathological hallmarks. We also find that several centenarians have no or only low amounts of Aβ pathology and intermediate levels of NFT loads, corresponding with the neuropathological description of primary age-related tauopathy (PART), [9]. The relation between PART and the occurrence of dementia is still elusive and its implications for cognition are still debated [11, 18]. The centenarians with PART in this cohort show variable performance on neuropsychological tests.

Thus far, many neuropathological changes such as ARTAG, atherosclerosis, hippocampal sclerosis, pTDP-43, GVD and Lewy bodies, are reported to accumulate with age independently of cognitive decline [1, 20, 21, 27, 38]. All centenarians presented high stages of GVD pathology, atherosclerosis as well as ARTAG. In contrast, the accumulation of pTDP-43 was limited in centenarians and we observed no association with cognitive test performance [26, 38, 47]. pTDP-43 aggregation has been suggested to be facilitated by Aβ pathology [38], for which we find only weak evidence in centenarians. We find that higher pTDP-43 stages occurred in all cases with hippocampal sclerosis, which is in agreement with previous observations [25]. Hippocampal sclerosis has previously been associated with lower performance on neuropsychological tests [27], and here we find a significant association with the inability to complete tests, a possible proxy for fatigue. Likewise, the accumulation of α-synuclein (Lewy bodies) was also limited in centenarians, despite the reported increased occurrence of this pathology with age[38]. We did not observe a correlation between the presence of Lewy bodies and cognitive function. The overall low occurrence of TDP-43 and Lewy body pathologies in this group of centenarians implies that these pathologies are more likely disease-related and not aging-related. In addition, we found one odd case with a Braak stage 6 for Lewy bodies, who was diagnosed with Parkinson’s disease many years before death, but performed well on the MMSE.

In general, in our cohort of centenarians increased levels of pathology correlate with lower cognitive performance. This observation is in agreement with the idea that simultaneous accumulation of different neuropathological hallmarks of AD could collectively account for the cognitive decline that is associated with aging [18, 19, 28, 38]. It remains elusive whether these centenarians retained their cognitive health to extreme ages because they delayed the accumulation of pathologies to much later ages, or because build-up of pathology was slower.

Thus far, we present the findings in a sample of 40 centenarians. With the continuation of this prospective study, we will be able to validate these findings in a larger sample in the near future. Also, as of yet we did not observe a cognitively healthy centenarian who was free of atherosclerosis or AD-associated pathologies, as reported previously for a 115 year old woman [10]. In line with reports that cognitive performance is highly correlated with survival in the oldest old [37], we find that the centenarians who score highest on cognitive tests in the ongoing prospective study are underrepresented among the first centenarians in the 100-plus Study who came to autopsy (data no shown). We therefore speculate that individuals with a low burden of pathology might survive until extreme ages, and will come to autopsy at a later time after inclusion.

In conclusion, we find that AD associated neuropathology is common in centenarians, albeit within limits for specific neuropathological hallmarks. Furthermore, we find that although the centenarians in this cohort escaped or delayed cognitive impairment until extreme ages, the accumulation of overall pathologies relate with lower cognitive performance. It remains unclear whether pathology accumulation was slower compared to others or whether accumulation started later. Overall, the registration of ante-mortem cognitive performance combined with thorough post-mortem brain analysis provides a unique window of opportunity for the association of cognitive function with the occurrence of neuropathological changes in centenarians.

## Acknowledgements

We thank and acknowledge all participating centenarians and their family members and the persons who contributed to this work as part of their bachelors’ and masters’ thesis: Neline van Woerden, Elleke Tissink, Melissa de Reus, Rianne Koeling, Ramon Bettings and Julia Letschert.

